# *Brucella* Peptide Cross-Reactive MHC I Presentation Activates SIINFEKL-Specific TCR Expressing T Cells

**DOI:** 10.1101/301465

**Authors:** Jerome S. Harms, Mike Khan, Cherisse Hall, Gary A. Splitter, E. Jane Homan, Robert D. Bremel, Judith A. Smith

## Abstract

*Brucella spp* are intracellular pathogenic bacteria remarkable in their ability to escape immune surveillance and therefore inflict a state of chronic disease within the host. To enable further immune response studies, *Brucella* were engineered to express the well characterized chicken ovalbumin (OVA). Surprisingly, we found that CD8 T cells bearing T cell receptors (TCR) nominally specific for the OVA peptide SIINFEKL (OT-1) reacted to parental *Brucella*-infected targets as well as OVA-expressing *Brucella* variants in cytotoxicity assays. Furthermore, splenocytes from *Brucella* immunized mice produced IFN-γ and exhibited cytotoxicity in response to SIINFEKL-pulsed target cells. To determine if the SIINFEKL-reactive OT-1 TCR could be cross-reacting to *Brucella* peptides, we searched the *Brucella* proteome using an algorithm to generate a list of near-neighbor nonamer peptides that would bind to H2K^b^. Selecting five *Brucella* peptide candidates, along with controls, we verified that several of these peptides mimicked SIINFEKL resulting in T cell activation through the “SIINFEKL-specific” TCR. Activation was dependent on peptide concentration as well as sequence. Our results underscore the complexity and ubiquity of cross-reactivity in T cell recognition. This cross-reactivity may enable microbes such as *Brucella* to escape immune surveillance by presenting peptides similar to the host, and may also lead to the activation of autoreactive T cells.

## INTRODUCTION

Brucellosis is a zoonotic disease caused by the gram-negative, facultative coccobacilli bacteria of the genus, *Brucella*. *Brucella* spp reside intracellularly within the host organism, preferring macrophages and macrophage-related cells. However, they also can persist extracellularly or outside the host. Symptoms of the disease are variable, including undulant fever, osteoarticular, genitourinary, and neurological complications. Within the host, *Brucella* have demonstrated the ability either to hide from or misdirect the immune response leading to chronic disease and complicating vaccine development (1). Although cytotoxic T lymphocytes (CTL) are a potentially major contributor to the control of brucellosis (2–4), the actual role of MHC class I-restricted CTL is unclear. One study demonstrated that the absence of perforin did not affect the level of infection (5, 6). On the other hand, in the study by Oliveira et al. β2m−/− mice were impaired in containment of Brucella infection(7), and Murphy et al. showed that CD8 T cell depletion exacerbated disease (8). *Brucella* have the ability to sabotage adaptive immune response, through undefined suppressive or regulatory means leading to the appearance of apparently exhausted CD8 T cells (3). The events producing exhaustion, as well as the nature of this state during chronic *Brucella* infection await better definition but nevertheless suggest that CTL could be key in limiting infection if not suppressed. In other model systems of CD8 exhaustion, notably lymphocytic choriomeningitis virus (LCMV), the study of T cell responses has benefited tremendously from the availability of specific research tools such as T cell receptor (TCR) transgenics. In comparison, *Brucella* research has been relatively hindered by the inability to identify antigen specific T cells. Although peptide epitopes have been published, there are no functional tetramers. To address this deficit, we sought to engineer *Brucella* to express a defined antigen that the infected antigen presenting cell (APC) would present in the context of MHC class I (MHC I) to more readily characterize the immune response to *Brucella* infection using a mouse model.

Due to its long history in immunological research, OVA is one of the best characterized model antigens, with epitopes that have been mapped for several mouse strains. Transgenic mice expressing the variable region of the TCR specific to the OVA peptide SIINFEKL (9) are referred to as OT-1. Every CD8+ T cell expresses this TCR transgene (10). The combination of OT-1-TCR-transgenic T cells and OVA-derived peptide SIINFEKL in the context of H2K^b^ is the most widely examined TCR-pMHC (peptide-MHC) complex (10, 11). Because of these readily available research tools, OVA has been a reference protein used to study CD8 T cell responses in other intracellular infections. Previous research has shown that intracellular bacteria such as *Listeria monocytogenes* or *Mycobacterium bovis* (BCG) expressing the OVA antigen induce strong antigen-specific primary and memory CD8 T cell responses (12–15).

In this study, we engineered and characterized OVA-expressing *Brucella* with the intent of studying primary and secondary CD8 T cell responses in acute and chronic brucellosis using the mouse model. Unexpectedly, we found the research tools used to analyze OVA antigen—specifically the cloned OT-1 TCR that recognizes SIINFEKL peptide presented by H2K^b^ —reacted to native *Brucella* infection as well. We therefore hypothesized that the *Brucella* proteome contains sequences similar to, or mimicking, the OVA SIINFEKL peptide. These results suggest the OT-1 TCR transgenic mice may be used to study native *Brucella* infections and further raises questions about the nature of cross presentation and molecular mimicry.

## MATERIALS AND METHODS

### Mice

C57BL/6 (Harlan) and C57BL/6-Tg(TcraTcrb)100Mjb/J (Jackson) were housed and cared for in AAALAC certified facilities of the University of Wisconsin School of Veterinary Medicine. Care, handling and experimental procedures were approved by the Institutional Animal Care and Use Committee (IACUC) with strict adherence.

### Cells and Cell culture

*Brucella melitensis* 16M strains and all *Escherichia coli* strains used in this project were cultured in Brain Heart Infusion (BHI) broth or agar at 37°C. Mouse dendritic cell line DC 2.4 (H2K^b^), and mouse monocyte cell line LADMAC were cultured in RPMI supplemented with 10% FCS, and 1mM sodium pyruvate (R10) in a humidified 37°C incubator with 5% CO_2_. The B3Z CD8+ T cell hybridoma cell line, specific for the SIINFEKL (OVA_257–264_/K^b^) peptide of OVA, was a kind gift from Dr. J.D. Sauer (University of Wisconsin-Madison). B3Z cells were cultured in R10 + 500 μg/ml G418 (Geneticin). Bone marrow derived macrophages (BMDM) were prepared by the culturing of bone marrow cells from the tibia/fibula of mice in R10 conditioned with 20% LADMAC supernatant.

### Plasmid and transposon engineering

We used the EZ-Tn5^™^ (Lucigen) transposon mutagenesis system for random insertion into the *Brucella* genome following the manufacturer’s recommended protocol. The insert was cloned into the transposon construction vector pMOD^™^-3 <R6Kγ*ori*/MCS> so that rescue cloning could be performed to determine the insertion site within the transformed *Brucella.* The partial OVA sequence was amplified from the vector pPL2erm-ActA100-B8R-OVA (kind gift from Dr. J.D. Sauer). Primers incorporated a *Brucella* ribosome binding site (RBS) designed using the algorithm, RBS Calculator v2.0, for high translation initiation (16, 17). The primers also contained *Eco*RI and *Bam*HI Restriction sites for subcloning into pECFP-N1 (Clontech) to produce an OVA-CFP fusion protein. Primers were as follows: N’-TGAAAGCAAAAGCAGAGAATTCTGGAATATTTTAATTCAGTATCAAAGAGAGGTAAACATGCAAGCCAGAGAGCTCATCA; C’-TTGAGGATCCTTCAGGCTCTCTGCTGAGGAGATGCCAGACAGA. PCR was performed using GoTaq^®^ Flexi DNA Polymerase system (Promega) with 6 mM MgCl_2_, and 55°C anneal temperature. The 632 bp product was subcloned into pECFP-N1. The fusion product was inserted into the *Eco*RI/*Xba*I site of pMOD-3. Finally, the Kanamycin resistance sequence was added to the *Sal*I site from pUC4k (Amersham) that we had modified to be flanked by loxP sites. The final product was named pMOD3-OVA-CFP. The map can be seen in **supplemental Figure S1**.

### *Brucella* transformation and rescue cloning

Transposons were generated by PCR following the manufacturer’s recommended protocol (Lucigen). Electrocompetent *Brucella* were prepared by growing *Brucella* to log phase in BHI broth. *Brucella* was pelleted and washed at least four times with ice cold water. The electrocompetent *Brucella* (50 μl) was then electroporated with the transposon (2 μl). Then, 950 μl of BHI was immediately added to the cells followed by overnight shaking in an incubator at 37°C. The next day, 200 μl of cells were plated on BHI agar plates containing 50 mg/L kanamycin. Plates were cultured for 5-7 days at 37°C. Clones were then selected and cultured in 96-well plates as a bacterial library and clones from the library were then propagated for rescue cloning of the transposon insertion site. Rescue cloning was performed following the manufacturer’s (Lucigen) protocol. Briefly, *Brucella* transformant genomic DNA was extracted using MasterPure DNA purification kit (Epicentre) and 2 μg of DNA was digested to completion with *Nco*I overnight to generate a fragment with intact transposon and flanking sequences. Digested DNA was religated using a FastLink DNA ligation kit (Epicentre). Ligations were column purified and transformed into electrocompetent EC100D*pir*+ cells (Epicentre) and plated on BHI agar containing kanamycin (50 μg/ml). Kanamycin-resistant colonies were selected, the plasmid was extracted, and the site of insertion was identified by sequencing the plasmid DNA bidirectionally using outward primers [forward (FSP; 5’-GCCAACGACTACGCACTAGCCAAC) and reverse (RSP; 5’-GAGCCAATATGCGAGAACACCCGAGAA)]. Sequencing was performed at the DNA sequencing core facility of the University of Wisconsin Biotechnology Center. Sequences were compared to the 16M genome sequence to determine the site of insertion.

### Western Blotting

Protein lysate of both bacteria and mammalian cells extract was made using B-PER^™^ Bacterial protein extraction reagent (ThermoFisher). Proteins were prepared for SDS-PAGE by heat denaturation in Laemmli sample buffer (BIORAD). Equal amounts of protein were added to wells of a 4-20% Tris-HCl gradient gel (BIO-RAD) along with SuperSignal^®^ molecular weight protein ladder. Separated proteins were transferred to nitrocellulose (BIO-RAD). Western blotting was performed utilizing a Pierce^®^ Fast Western blot kit following the manufacturer’s instructions. Antibodies included mouse monoclonal anti-OVA (3G2E1D9; GenTex), and mouse monoclonal anti-GFP (B-2; SCBT) used at recommended dilutions. Chemiluminescent blots were visualized on X-ray film.

### Fluorescence Microscopy

BMDM were prepared from C57BL/6 mice and plated on chambered coverslips (IBIDI). Some samples were then infected (1000 MOI) with *Brucella* expressing tdTomato fluorescent protein (CLONTECH) for 24 h. The high MOI was chosen to increase sensitivity consistent with our previous studies (18). Other samples were pulsed with SIINFEKL peptide (50 μM) Cell samples were fixed in 4% paraformaldehyde and processed for fluorescence confocal microscopy. Cells were stained with monoclonal antibody to OVA 257-264 (SIINFEKL) peptide bound to H2K^b^ (eBioscience) and then with goat anti-mouse IgG (H+L) Alexa Fluor 488 (Dylight; ThermoFisher). Imaging was performed at the University of Wisconsin Optical Imaging Core using either a Nikon A1RS confocal microscope or a Leica SP8 3X STED Super-resolution microscope.

### Cytotoxic T Lymphocyte (CTL) Assay

For the CTL assays, assessment consisted of measuring extracellular activity of dead-cell protease by luminescence using the CytoTox-Glo^™^ Cytotoxicity Assay (Promega). Effector and targets for the assay varied as described below. Immune effectors were prepared as follows: Mice (C57BL/6, female, 4wks old) were injected intraperitoneally with PBS (diluent control) or 2 × 10^6^ *Brucella* in 200 μl PBS (*B. melitensis*, *B. melitensis* ova-cfp #3, *B. melitensis* ova-cfp #4). Each group consisted of 4 mice. After three weeks, mice were euthanized and splenocytes were harvested. CD8+ T cell effectors were isolated from splenocytes using a MACS CD8a+ T cell isolation kit (Miltenyi). Targets consisted of splenocytes from age-controlled mice that were pulsed with OVA_257-264_ peptide (SIINFEKL) or not peptide-pulsed. Briefly, mononuclear splenocytes were suspended in complete growth media (R10) at 5 × 10^6^/ml. OVA peptide (SIINFEKL; GenScript) was added at 1 μl/ml from a 200 μM stock and cells were incubated at 37°C for 1h. To control for non-specific cytotoxicity, unpulsed target controls were also included in the assay and this background level was subtracted from the experimental levels. Targets, effectors, and controls were plated in triplicate in 96-well round bottom plates. Cells were incubated for 5 h then assayed by luminometry. Specific cytotoxicity represents SIINFEKL pulsed target cell death minus background non-pulsed target cell death. The OT-1 effector cytotoxicity assay was prepared as follows: Splenocytes from OT-1 mice (C57BL/6-Tg(TcraTcrb)1100Mjb/J) were processed and CD8+ T cell effectors (OT-1 cells) were isolated using a MACS CD8a+ T cell isolation kit (Miltenyi). Targets consisted of DC2.4 mouse (H2K^b^) dendritic cell line that were either non-infected (control), or infected (MOI 100) overnight with *Brucella* (*B. melitensis*, *B. melitensis* ova-cfp #3, *B. melitensis* ova-cfp #4). Positive control were cells pulsed with OVA peptide (SIINFEKL) as described above. Targets, effectors, and controls were plated, incubated, and assayed as described for immune effectors CTL assay above following the manufacturer’s (Promega) recommended protocol and calculations for percent specific cytotoxicity.

### IFN-γ Enzyme Linked Immunosorbent Assay (ELISA)

Effector cells were prepared by immunizing mice (C57/BL6) as described for the CTL experimental group above except that an additional group was immunized with OVA peptide (SIINFEKL) at 50 μg in 0.2 ml Sigma Adjuvant System^®^ (Sigma) i.p. following the manufacturer’s protocol. Peptide immunizations were boosted after 2 weeks and one week later, splenocytes were harvested. Splenocytes from each animal were then stimulated in culture with 1 μg/ml of SIINFEKL peptide and incubated for 48 h at 37 °C. Cultured supernatants were harvested, and IFN-γ amounts were determined using the ELISA Ready-Set-Go! System (Affymatrix; eBioscience) following the manufacturer’s protocol.

### β-Galactosidase X-Gal and ONPG Assays

Cultures containing a mix of B3Z T cell hybrids and DC2.4 APCs (2 × 10^5^ cells/ml each were plated in 6-well tissue culture plates (X-Gal assays) or 96-well flat bottom tissue culture plates (ONPG assays) and peptide or bacteria added. Peptides (listed in **Table 2**) were synthesized and purchased from GenScript and resuspended in DMSO at a stock concentration of 20 mg/ml. *B. melitensis* and variants were used at 100 MOI. For some assays, APCs were treated with Tauroursodeoxycholic acid (TUDCA; Sigma) at 100 μg/ml, or Tunicamycin (Sigma) at 10μg/ml, or mouse IFN-γ (PromoKine) at 1 μg/ml. After overnight incubation, cells were washed in PBS and fixed (X-Gal assay) or lysed (ONPG assay). For X-Gal staining, cells were fixed with 4% PFA for 10 min, then washed 3X in PBS, and overlaid with a solution of 1 mg X-Gal/ml, 5 mM potassium ferrocyanide, 5 mM potassium ferricyanide, and 2 mM MgCl_2_. After an overnight incubation at 37°C, plates were examined microscopically for the presence of blue (lacZ expressing) cells. For ONPG staining, we used a SensoLyte^®^ ONPG β-Galactosidase Assay kit (AnaSpec, Inc) following the manufacturer’s recommended protocol except that the incubation at 37°C was overnight. Absorbance reading was at 405 nm.

### Prediction of near neighbors

Structural diagrams of binding of SIINFEKL in the H2K^b^ murine MHC I molecule (3P9L) (19) illustrate that this octomer is bound with its C terminal leucine located in the P9 pocket position and the N terminal serine in the P2 pocket position. This results in a pentamer peptide exposed to the T cell receptors (T cell exposed motif or TCEM) (20). Amino acids N, E and K protrude particularly prominently from the MHC groove in positions P5, P7, and P8. By replacing amino acids with those of similar physicochemical characteristics in T cell exposed positions P4-P8, 31 peptides were identified which a T cell receptor would likely tolerate and bind as an alternate “near neighbor” of SIINFEKL. The amino acid substitutions in the exposed positions included P4: I>L, P5: N>Q, P6: F>I, A, L or Y, P7: E>D, P8: K>R. The *B. melitensis* proteome and the proteomes of an array of other pathogenic and microbiome bacteria were then searched to determine the occurrence of each of the alternate TCEM pentamer motifs P4-P8. Where a near neighbor was identified in *B. melitensis*, the flanking amino acids were noted, and the predicted binding to murine MHC I alleles was determined in the context of the native *Brucella* protein using previously described methods (21), and the probability of cathepsin cleavage at the C terminus of that peptide determined (22, 23).

### Statistical Analyses

Analysis of variance (ANOVA) was used to analyze the differences among group means. Tukey’s HSD (honest significant difference) test was used as the post hoc follow-up test comparing every group mean with every other group mean to determine significant differences among groups.

## RESULTS

### Engineering and characterization of OVA antigen expressing *Brucella*

Our long-term objective being acute and chronic brucellosis immunological studies, we engineered *Brucella* to express well-characterized antigens with readily available antigen-specific research tools. *Brucella melitensis* 16M was transformed to express a fusion protein consisting of a fragment of chicken ovalbumin (OVA) and Cyan Fluorescent Protein (CFP). This fusion protein sequence was determined to have a predicted probability of antigenicity of 0.9 as measured by ANTIGENpro software using the SCRATCH protein predictor (http://www.ics.uci.edu/~baldig/scratch/index.html).

The nucleic acid sequence contains a Ribosome Binding Sequence (RBS) optimized for *Brucella* and the promoter would be provided by the insertion gene (Figure 1). The OVA sequence selected contained OVA_257-264_ (SIINFEKL); the well-characterized H2K^b^ restricted peptide epitope (24) and the CFP portion contained the H2K^d^ restricted epitope HYLSTQSAL (25). A library of *Brucella* transposon transformants was made and rescue cloning performed to determine the transposon insertion site (**Table 1**). Western blotting of chosen clones using anti-OVA or anti-GFP specific antibodies determined protein expression (Figure 2). Viability of transformed clones were compared to parental *Brucella* by growth in broth as well as intracellular growth *in vitro* and *in vivo* measuring CFU from infected BMDM in culture or CFU from splenocytes of infected mice (Figure 3). Clones with comparable growth to wild type (clones #3 and #4) with insertion in *BMEI 1025* and *BMEII 0058* respectively (Table 1), were chosen for further evaluation.

**Figure 1.**
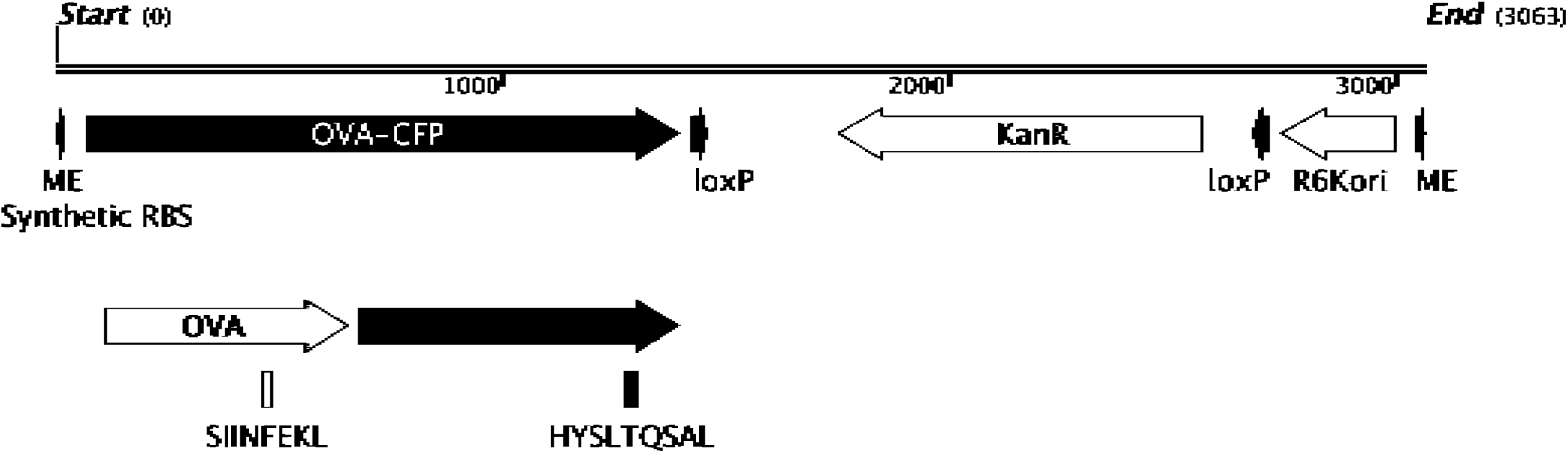
Transposon map of inserted elements. OVA-CFP and kanamycin coding sequence along with synthetic RBS and R6K origin of replication for rescue cloning are displayed. Also represented are the locations of SIINFEKL peptide in the partial chicken ovalbumin sequence and HYSLTQSAL peptide in the ECFP sequence. ME (mosaic ends) and loxP sites for Cre/lox recombination are also displayed.

**Figure 2.**
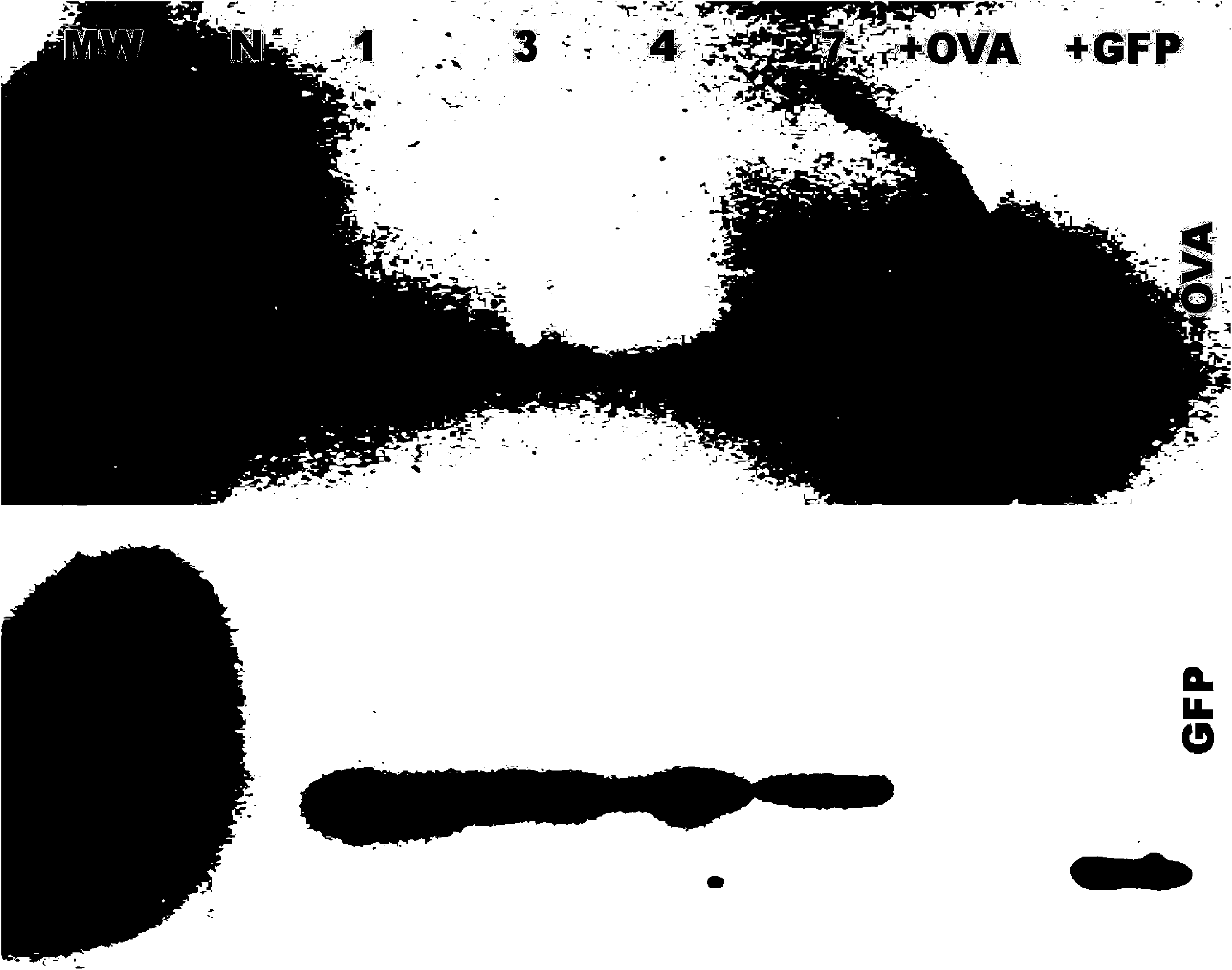
Western analyses of Transposon transformed *Brucella* lysates. Equivalent amounts of *Brucella* clones were lysed, denatured in SDS-laemmli sample buffer, PAGE separated, and blotted. Antibodies to chicken ovalbumin (OVA) or Green Fluorescent Protein (GFP) were used along with HRP secondary antibody. Chemiluminescence was detected by X-ray film. (N) parental *Brucella*; (1, 3, 4, 7) Transposon Transformed *Brucella* clones; (+OVA) chicken ovalbumin; (+GFP) Green Fluorescent Protein; (MW) molecular weight marker.

**Figure 3.**
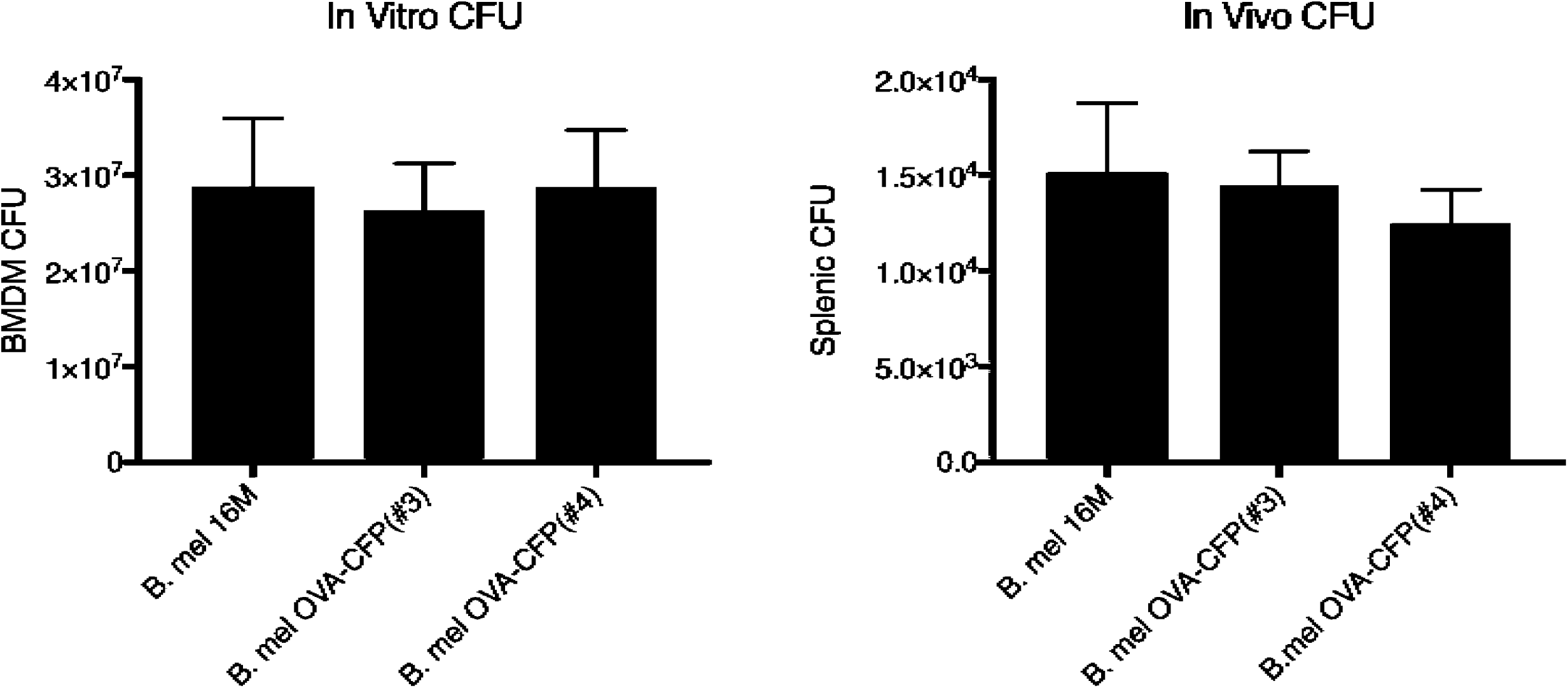
*In Vitro* and *In Vivo* Colony Forming Units (CFU) of *Brucella* infected macrophages and splenocytes. Parental *Brucella* along with transformed *Brucella* clones #3 and #4 were used to infect C57BL/6 Bone Marrow Derived Macrophages (BMDM) in culture (100 MOI) for 24 h or infect mice (C57BL/6) at 2 × 10^6^ bacteria for 7 days. Data are representative of three experiments.

### OVA_257-264_ (SIINFEKL) is presented by H2K^b^ in OVA-expressing *Brucella* infected mouse BMDM

The next objective was to determine whether the OVA-GFP fusion protein expressed by the *Brucella* transformants could be processed and presented by host cell MHC I. More specifically, to determine if the OVA SIINFEKL peptide would be processed and presented on cell surface MHC class I, we employed an antibody specific for H2K^b^ bound to SIINFEKL peptide. BMDM from C57BL/6 mice were infected with OVA-expressing *Brucella* and analyzed by fluorescence microscopy (Figure 4). Results indicate that SIINFEKL peptide-MHC I complexes could be directly visualized on the *Brucella*-OVA infected macrophages.

**Figure 4.**
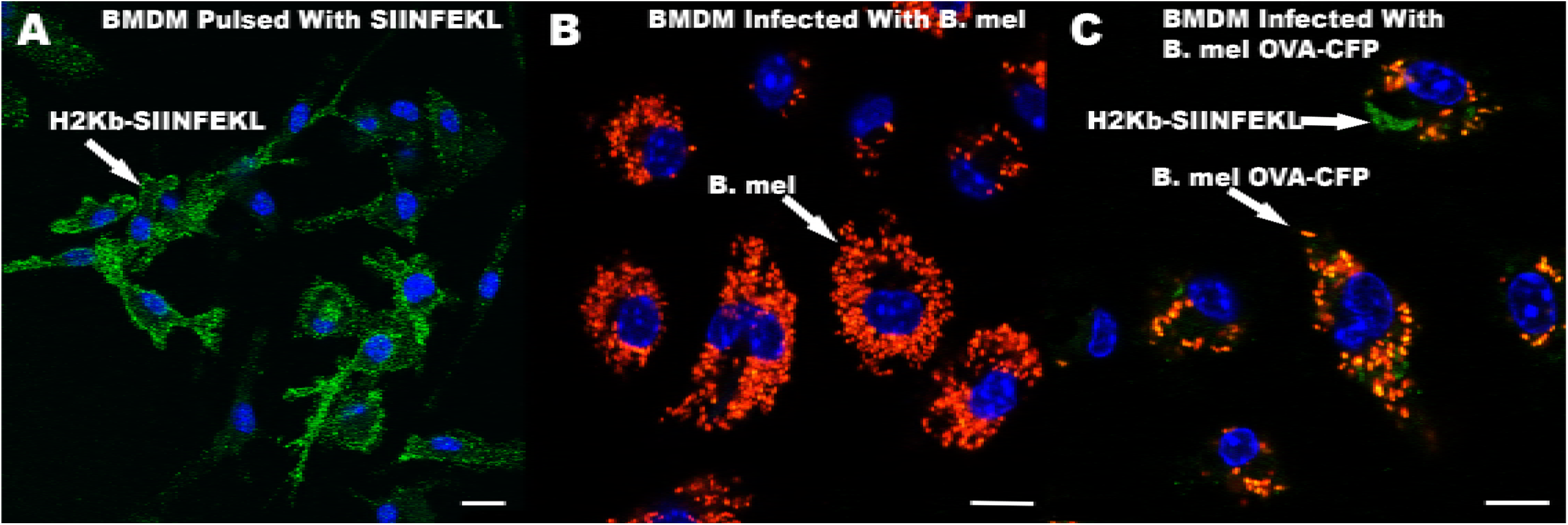
Fluorescence microscopy analyses of H2K^b^-SIINFEKL. BMDM from C57BL/6 mice were either pulsed with SIINFEKL peptide (A), infected with 1000 MOI parental *B. melitensis* (B. mel) (B), or infected with 1000 MOI *B. melitensis* OVA-CFP (clone #4)(C). Both *B. melitensis* strains expressed tdTomato (Red). Cells were fixed and stained with H2K^b^ -SIINFEKL antibody (green). SIINFEKL pulsed cells indicated H2K^b^ was bound to SIINFEKL as expected. Scale bar represents 10 μm.

### Antigen from wild type *Brucella* mediates OT-1 (OVA-specific) T cell activation and generates effectors capable of recognizing SIINFEKL bound MHC I

As these results indicated that the OVA-CFP fusion protein was processed in infected cells, and that SIINFEKL was presented by H2K^b^, we proceeded to T cell immune response studies. Unexpectedly, CD8+ effector T cells from mice immunized with parental and OVA expressing *Brucella* both reacted to SIINFEKL-pulsed targets (Figure 5A). Indeed, effectors from control *B. melitensis* immunized animals lysed OVA peptide pulsed targets at a level not significantly different than the OVA-expressing *Brucella.* These data indicate that *B. melitensis* immune cells can recognize OVA peptide-MHC I.

**Figure 5.**
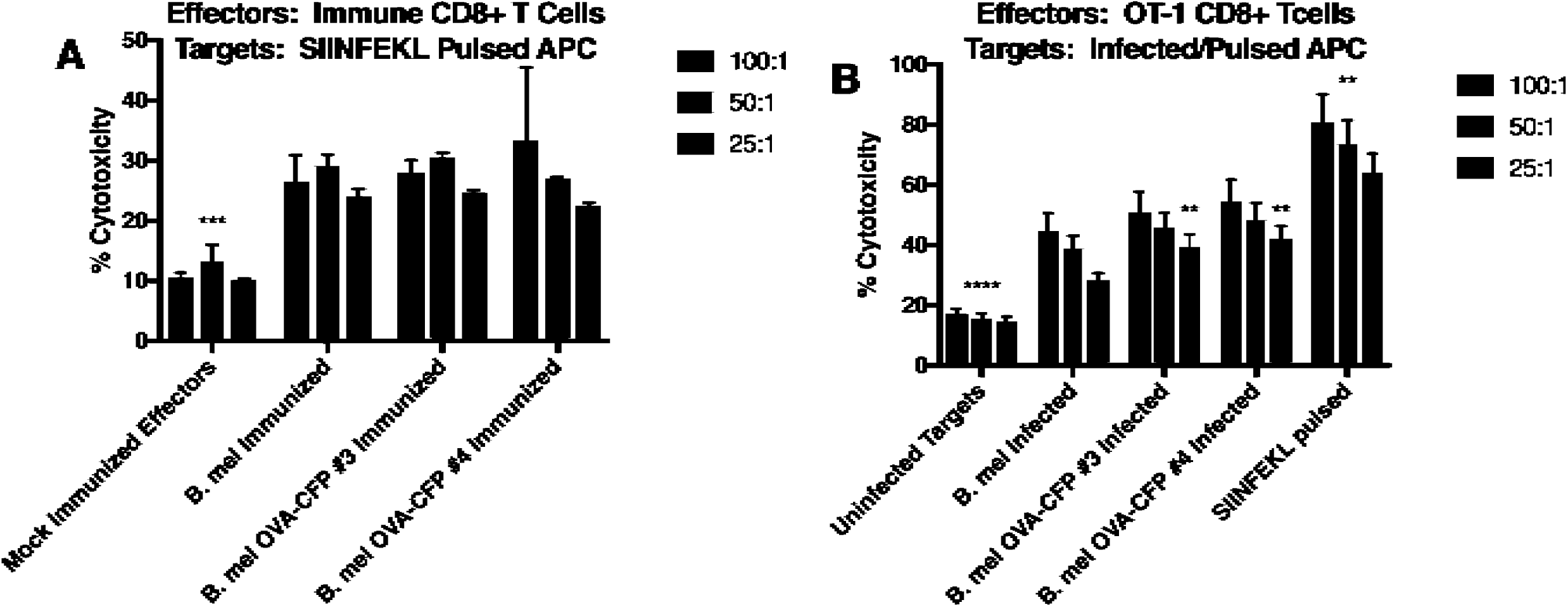
Cytotoxicity assays. In left panel (**A**), C57BL/6 mice were immunized with parental *B. melitensis* or OVA expressing variants. Effector splenic CD8+ cells from these mice were used against SIINFEKL-pulsed splenocyte targets at the indicated effector to target ratio (E:T). Non-specific cell death in unpulsed targets was subtracted to yield SIINFEKL specific cytotoxicity. In right panel (**B**), OT-1 CD8+ effectors were used against parental *B. melitensis* or OVA expressing variant infected target DC2.4 dendritic cells. SIINFEKL-pulsed targets were used as positive control. Data are representative of four independent experiments. ** p<.05; SIINFEKL-pulsed targets were significantly different from *B. melitensis* and variants at all E:T. Additionally, **p<.05; clones #3 and #4 were significantly different from parental *B. melitensis* at 25:1. ***p<.001; significantly different from immunized groups, ****p<.0001; significantly different from infected or peptide-pulsed targets. P-values reflect one way ANOVA statistical analyses.

Similar results were observed in separate experiments examining IFN-γ cytokine production. *B. melitensis* immunized effectors produced IFN-γ in response to SIINFEKL peptide at levels not significantly different from the *B. melitensis* OVA immunized effectors (Figure 6A). As expected, IFN-γ expression from *Brucella* immunized animals was significantly less than from OVA-peptide immunized animals. To confirm that the immunized cell response to SIINFEKL was specific, a separate immunization experiment was performed using SIINFEKL and a scrambled peptide to pulse the cells (Figure 6B). Production of IFN-γ by splenocytes from *Brucella*-immunized mice was also significantly higher in the presence of SIINFEKL peptide as compared to a scrambled control peptide.

**Figure 6.**
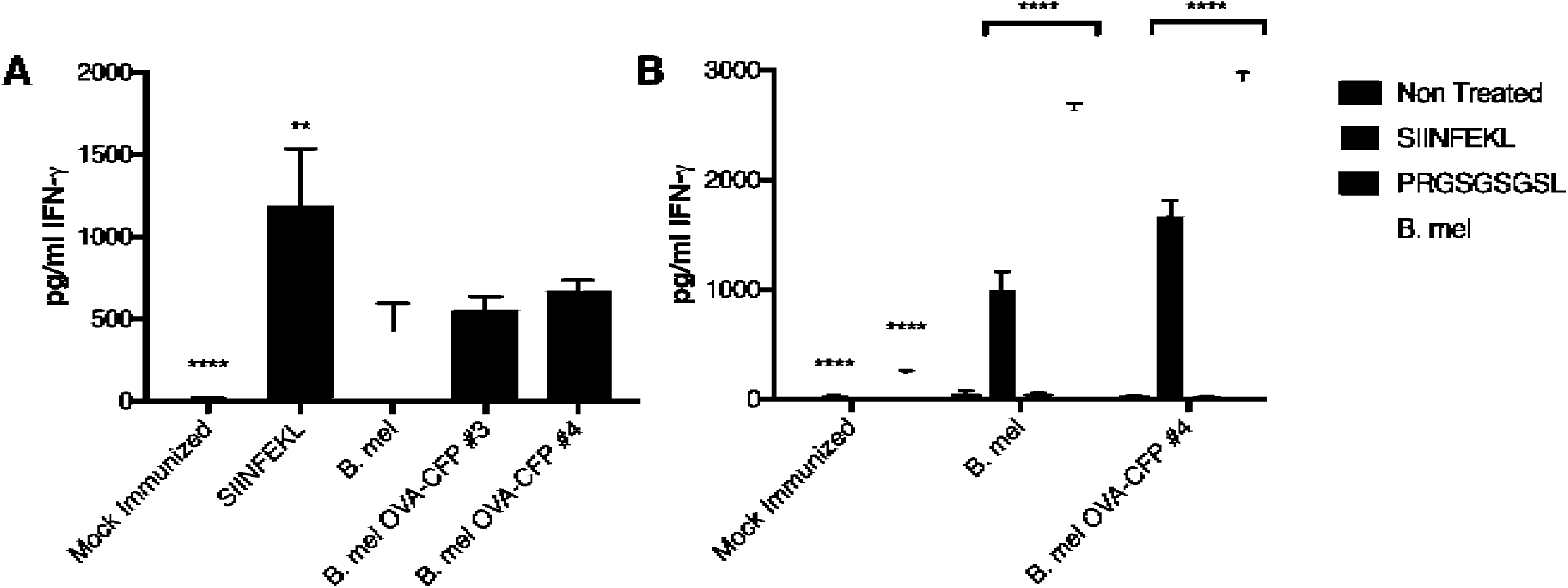
IFN-γ production by effectors from immunized mice. C57BL/6 mice were immunized with parental *B. melitensis* or OVA expressing variants (#3, #4), or SIINFEKL peptide in adjuvant as a positive control. In panel **A**, splenocytes were pulsed with SIINFEKL peptide and assayed for IFN-γ production by ELISA. In panel **B**, splenocytes from a separate experiment were either not pulsed (Non-Treated) or pulsed with SIINFEKL, PRGSGSGSL (random, negative control peptide), or infected with *B. mel* (MOI 100). **p<.05; statistically different from *B. melitensis* and variants. ****p<.0001; statistically different from peptide immunized or infected splenocytes. One-way ANOVA statistical analyses.

To affirm these unexpected results, we altered our CTL assay approach by utilizing the well characterized SIINFEKL-specific T cell receptor expressing OT-1 CD8+ T cells from TCR transgenic mice. This time the effectors (OT-1) were OVA-peptide specific and the targets were *Brucella* infected. Targets pulsed with SIINFEKL served as a positive control. Again, non-OVA expressing *Brucella* infected targets were lysed by OT-1 effectors at similar levels to OVA expressing *Brucella* (Figure 5B). As expected, OVA-peptide pulsed targets lysed at significantly higher levels.

One possible explanation of the results presented above could be that *Brucella* infection non-specifically activates T cells because of its effect on antigen presenting cells, or that we may be observing cross-reactivity of the OT-1 receptor to mouse peptides. *Brucella* infection induces ER stress and IFN production, either of which could potentially modulate antigen presentation (1, 18, 26). The SIINFEKL specific responses by *Brucella*-immune cells were significantly greater than un-pulsed or scrambled controls (Figure 6B), arguing against non-specific host stimulation of T cells as the sole explanation. However, to address this theory more directly, we treated the antigen presenting cells with infection-associate factors that could potentially alter MHC I-peptide presentation. The drugs TUDCA and Tunicamycin inhibit or enhance the Unfolded Protein Response (UPR) respectively (27). IFN-γ is known to enhance MHC I expression (28). To further isolate effects on antigen presentation and simplify responder population, we utilized the B3Z CD8+ T cell hybridoma with a TCR specific for the OVA (SIINFEKL)-H2K^b^ complex. The cell line was transfected with a lacZ reporter gene driven by the NFAT (nuclear factor of activated T cells) element of the human IL2 enhancer. The H2K^b^ presentation of SIINFEKL to B3Z cells activates NFAT and results in β-galactosidase synthesis, which can be detected as blue staining of cells visualized by microscopy or quantitated by development of the ONPG chromogenic substrate. The B3Z reporter system is widely used in T cell activation studies (29). Results shown in Figure 7 indicate that ER stress modulation or IFN-γ treatment did not result in further activation of B3Z reporter T cells bearing the OVA-specific TCR, by SIINFEKL peptide or whole *Brucella* infection.

**Figure 7.**
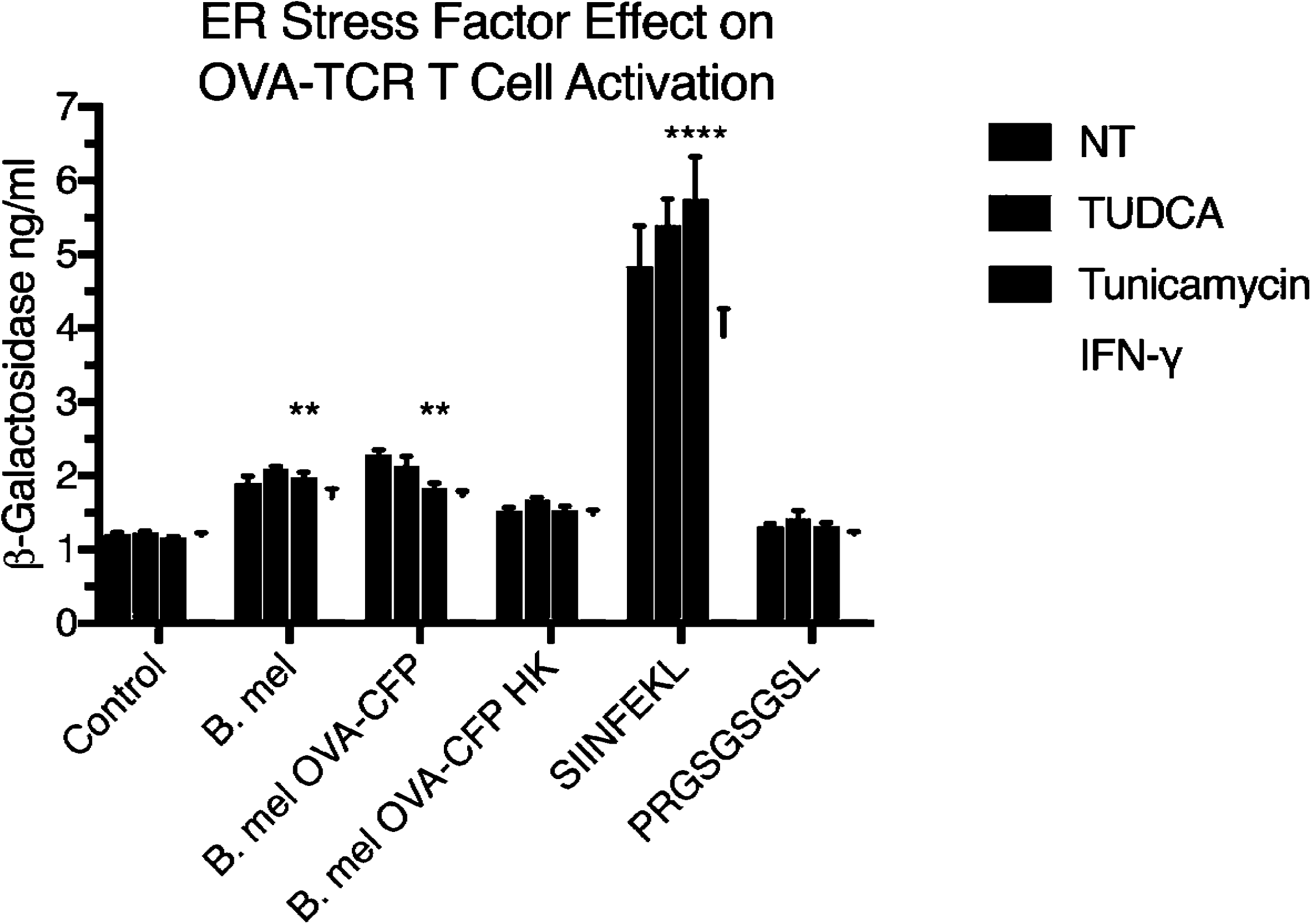
ER stress and IFN-γ effects on B3Z T cell hybrid activation by infected or peptide pulsed APCs. DC2.4 dendritic cells were not treated (NT) or treated with UPR inhibitor TUDCA (500 ng/ml), UPR inducer Tunicamycin (50 ng/ml) or IFN-γ (10 ng/ml) for 24h. APCs were subsequently infected (100 MOI) with *Brucella melitensis* (B. mel), OVA-expressing *B. mel* (*B. mel* OVA-GFP) or Heat-Killed OVA-expressing *B. mel* (B. mel OVA-GFP HK) or pulsed (10 μg/ml) with peptide for 24h. Control DC2.4 cells were not infected or pulsed with peptide (SIINFEKL or scrambled negative control). B3Z T cell hybrids were added and levels of TCR activation were measured through ONPG assay. Median and standard deviation of three experiments are shown. **p<.05, ****p<.0001; statistically different from control and scrambled peptide using one-way ANOVA statistical analyses.

### Detection of near neighbor T cell exposed motifs in *Brucella* indicates possible molecular mimicry

These surprising results led us to theorize that there might be cross-reactivity of the OVA-reactive T cells to structurally related peptides derived from *Brucella.* We performed a search of the *B. melitensis* proteome for the SIINFEKL sequence. The proteome of *B. melitensis* does not contain a peptide identical to SIINFEKL. However, 38 peptides were found that comprise a P4-P8 T cell exposed motif matching one of the peptides having near neighbor physicochemical characteristics. **Supplemental Table S1** shows these peptides, the *Brucella* proteins of origin and the predicted binding affinity to murine MHC I alleles H2K^b^ and H2D^b^ of the nonamer peptides that contain these motifs. For those peptides having the highest predicted binding affinity to H2K^b^ or H2D^b^, the probability of C terminal cathepsin excision was examined. Interestingly, for peptides which were predicted to have a high probability of C terminal cleavage by either cathepsin S or L to permit MHC I binding, cleavage probability was highest at the position P10, i.e. yielding a decamer peptide. This is consistent with prior observations that indicate that a decamer may be more likely to be initially excised than a nonamer (23). This selection process yielded a ranking of peptides for further study, where four were selected for testing. In making this selection we also considered proteins that had clearly detectable expression observed during our previous proteomics and RNAseq studies of infected cells (18). For controls we selected a peptide comprising a near neighbor motif that was predicted to have low affinity for H2K^b^ or H2D^b^, and a random nonamer peptide. **Table 2** describes the peptides used in these studies.

Examination of the proteomes of 4-5 distinct isolates each of 140 other bacterial pathogens, from 14 genera, identified that peptides comprising the P4-P8 motifs of near neighbors of SIINFEKL are not uncommon. The frequency of such near neighbor peptides in other pathogens are shown in **Supplemental Table S2**. Each bacterial proteome examined contains from 4 to 45 near neighbor motifs of SIINFEKL that may produce cross reactions similar to those shown here for *B. melitensis*, if the flanking amino acids in each context are conducive to cathepsin cleavage and to MHC I binding. A similar incidence of the near neighbor peptides was detected in proteomes of 20 bacteria found in the gastrointestinal microbiome (data not shown).

### Putative *Brucella* peptides can activate OVA-specific TCR bearing T cells

To assay cross-reactivity of the putative *Brucella* peptides, we employed the B3Z cell line (30) and peptide pulsed DC2.4 mouse dendritic cell line (H2K^b^) as APC. Visual scanning of the lacZ stained cells revealed that all the peptides tested in **Table 2**, except the random sequence, had some level of staining above background (no peptide added). In fact, three of the peptides (KSIINAERL, PQKINIDRT, and KNKINLDKL) were observed to have staining similar to SIINFEKL (Figure 8). We then repeated our assays using peptide dilutions to assess avidity of the pulsed peptide-MHC I and TCR interaction. The ONPG assays (Figure 9) verified our lacZ results that three of the experimental peptides (KSIINAERL, PQKINIDRT, and KNKINLDKL) had high levels of TCR (NFAT) activation. ONPG assays also revealed different avidities for the peptides. Figure 9B shows that whereas SIINFEKL avidity did not change much at all dilutions tested, the *Brucella* peptide avidity decreased with dilution. Notably, the native *Brucella* KSIINAERKL peptide did not differ significantly from the SIINFEKL peptide until 0.1μg/ml.

**Figure 8.**
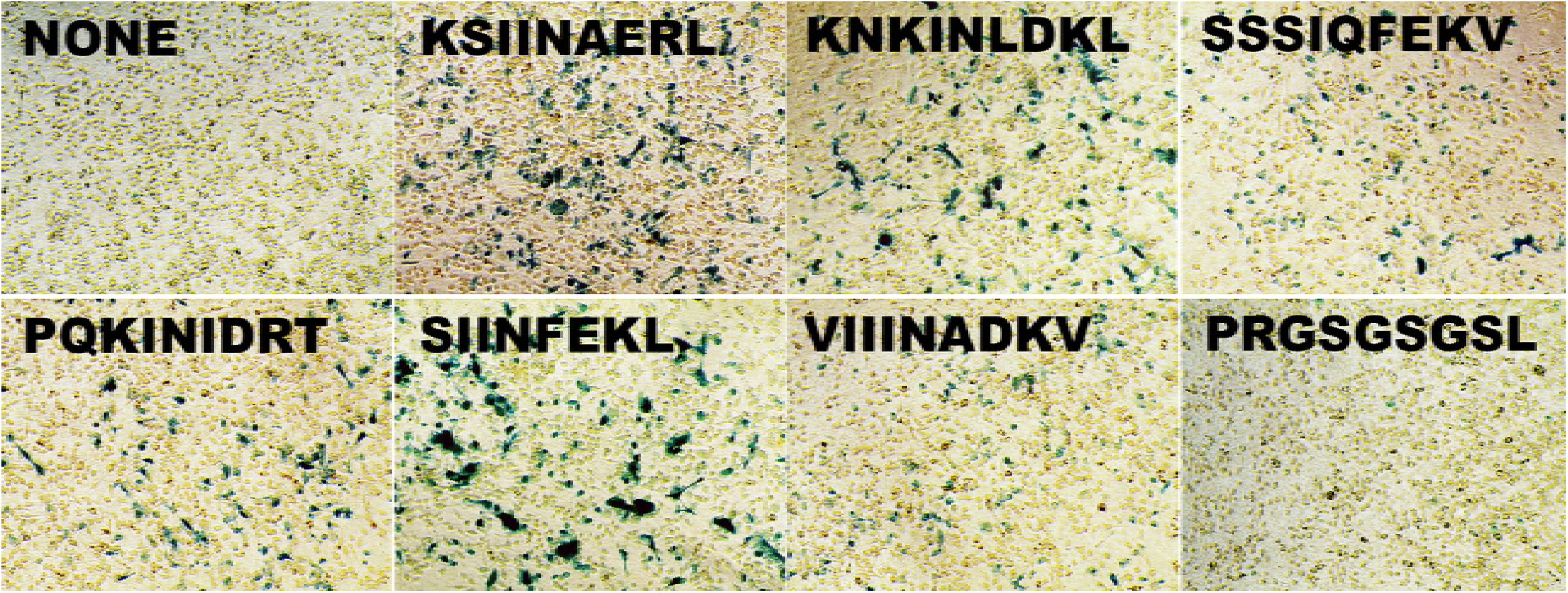
B3Z lacZ expression of T cell activation. DC2.4 dendritic cells were pulsed with 10 μg/ml of the indicated peptides peptides and mixed with B3Z T cell hybrids. PRGSGSGSL is the scrambled peptide control. After 24 h, T cell activation was monitored by β-galactosidase expression. Blue cells indicate peptide-MHC I binding of the TCR and T cell activation.

**Figure 9.**
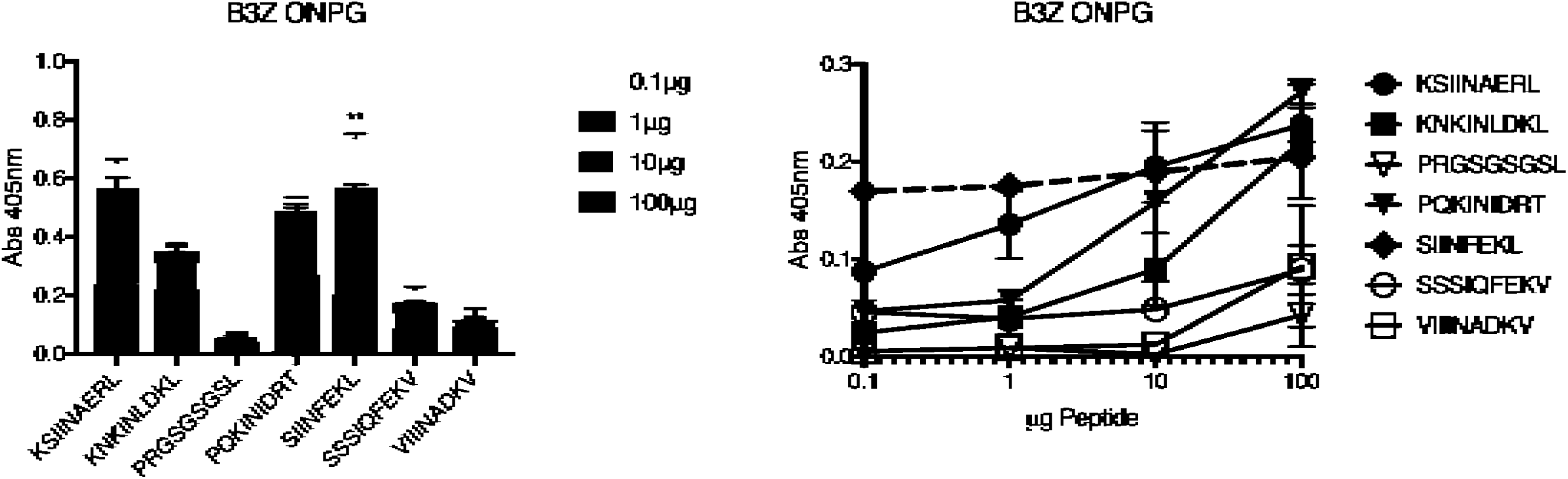
T cell activation by peptide-MHC I at different peptide dilutions. DC2.4 dendritic cells were pulsed with indicated amounts of peptide and mixed with effector B3Z T cell hybrids. Levels of TCR activation were measured through ONPG assay. Median and Standard Deviation of six assays are shown. Left graph (A) shows data expressed as stacked columns of various dilutions of each peptide. Right graph (B) shows the same data expressed as a line graph. Dashed line is SIINFEKL reference control. One-way ANOVA statistical analyses of median values indicated SIINFEKL treated samples were significantly different (**p<.05) from all other treatments at only 0.1 μg peptide concentration.

To determine if infection with the parental *B. melitensis* 16M generated immune responses to these OVA-TCR cross reactive peptides in vivo, splenocytes from mice immunized with either B. mel or B. mel-OVA were assayed for IFN-γ production after stimulation with peptide (Figure 10). Immune cells did indeed respond to the panel of cross-reactive peptides with greater cytokine production than to scrambled peptide, suggesting these epitopes may be generated *in vivo*.

**Figure 10.**
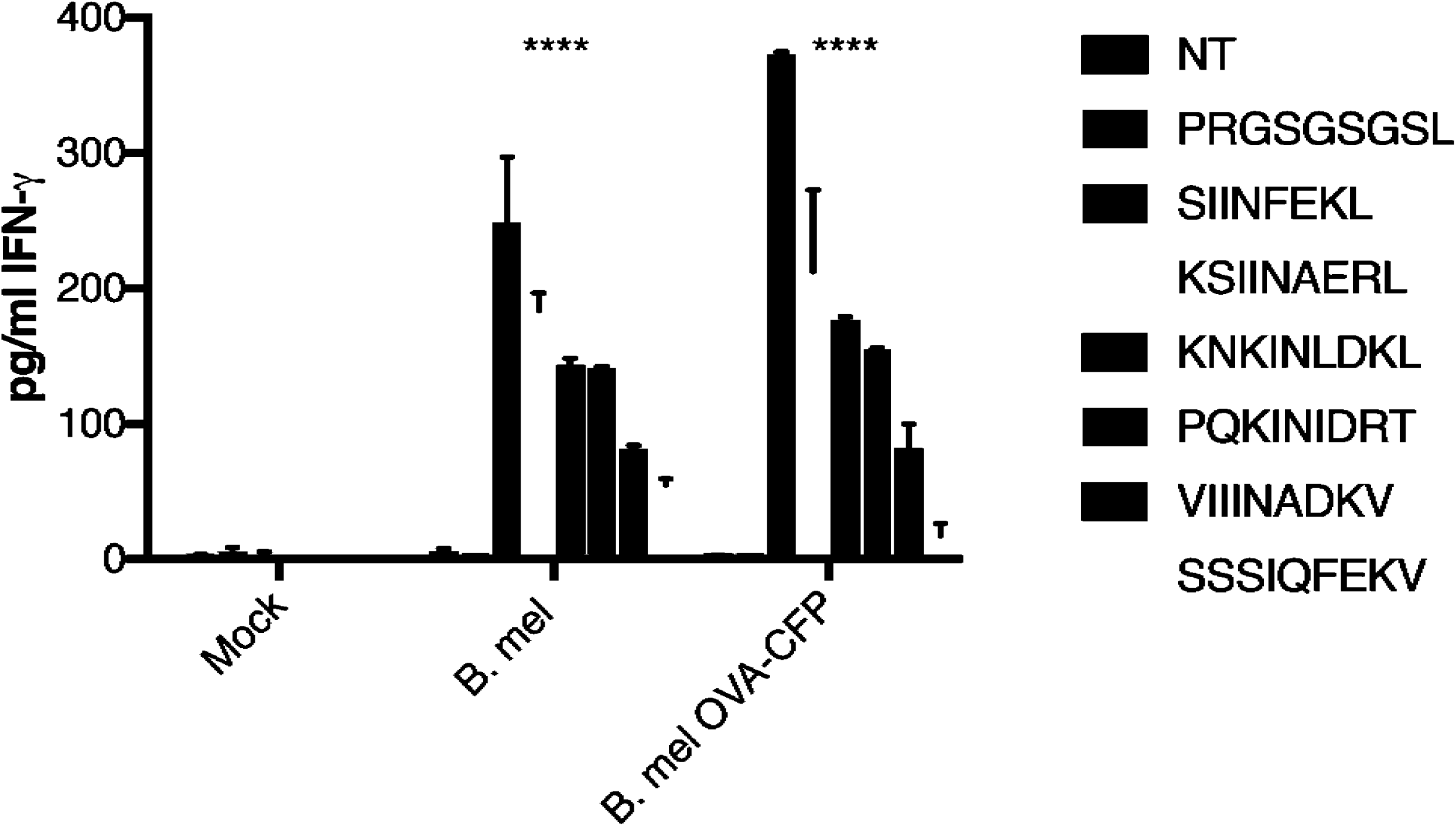
Immune response to peptides. C57/BL6 mice were immunized with 10^6^ *Brucella melitensis* (B. mel), *Brucella melitensis* expressing OVA-CFP antigen (B. mel OVA-CFP) or diluent (Mock). After 3 weeks, splenocytes were harvested and stimulated with peptide (50 μg) for 24 hrs and supernatant was assayed for IFN-γ by ELISA. Absorbance readings represent the median and standard deviation of four mice from each immunization group. ****p<.0001 significantly different from mock immunized group in two-way ANOVA statistical analyses.

Finally, we determined if these native *Brucella* peptides could activate OVA-specific TCR using the OT-1 mouse system. Splenocytes from these mice were pulsed with the various peptides and IFN-γ expression was measured by ELISA. Results shown in Figure 11 confirm our findings using the B3Z β-galactosidase reporter cell line that the OT-1 TCR is cross reactive to peptides of the *Brucella* proteome. Peptide immunization of the OT-1 mice was not attempted due to an anticipated hyper-immune response (9, 10). Indeed, when we tried infecting these mice with *Brucella* and the *Brucella*-OVA mutants, the mice died within six days (data not shown).

**Figure 11.**
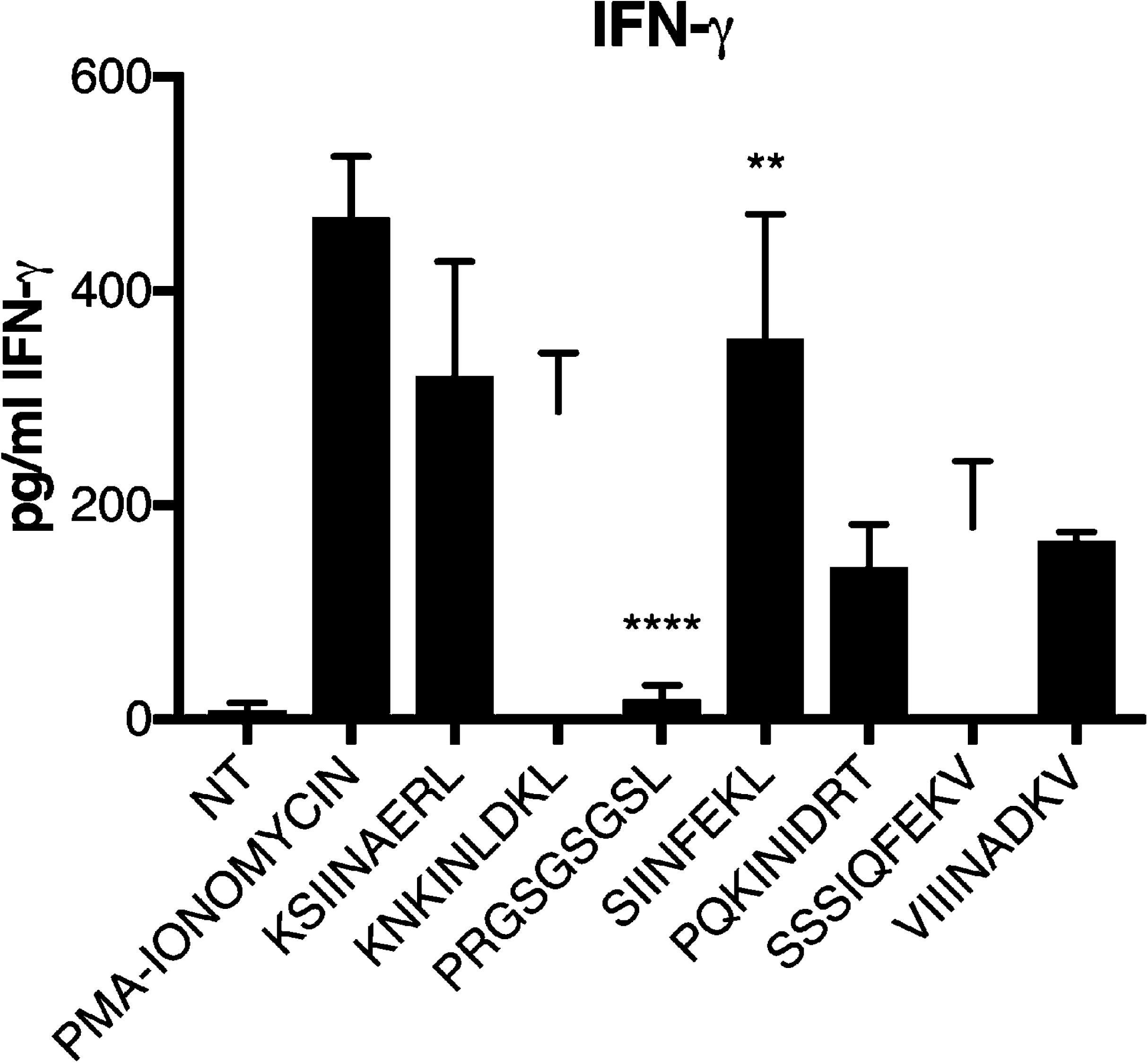
OT-1 CD8+ cell activation by native *Brucella* sequences. Splenocytes from OT-1 mice were pulsed with 50 μg of peptide for 24 hrs and supernatant was assayed for IFN-γ by ELISA. Absorbance readings represent the median and standard deviation of four experiments. NT; non-treated. **p<.05; SIINFEKL is significantly different from PRGSGSGSL, PQKINIDRT, SSSIQFEKV, and VIIINADKV. ****p<0001; PRGSGSGSL is significantly different from all other treated samples in one-way ANOVA statistical analyses.

## DISCUSSION

The immune response to *Brucella melitensis* is complex, as evidenced by the fact that there is currently no known effective brucellosis vaccine available. Our research goal was to engineer *Brucella* to express immunogenic OVA as a tool to follow antigen-specific effector and memory immune responses to *Brucella* infection *in vivo*, using the mouse model. Adopting this approach, we could employ SIINFEKL-MHC tetramers and T cells from TCR transgenic mice. This goal was nominally fulfilled. However, the surprising result was that native *Brucella* also stimulated “OVA-specific” TCR of OT-1 mice. Employing a panel of near-neighbor *Brucella* peptides, cross-reactivity of the OVA-TCR was evident, although to a lesser extent than the OVA SIINFEKL that was originally used in clonal selection of the TCR (10, 11, 31). Despite the lower avidity, as indicated by the B3Z assays, *Brucella* peptides from infections were sufficiently immunogenic to trigger both robust cytokine and CTL responses via the OVA TCR. These results suggest that the OT-1 TCR transgenic T cells may be used to probe native *Brucella* immune responses as well as responses to the *Brucella*-OVA.

A search of the *Brucella melitensis* proteome revealed several proteins containing T cell exposed pentamer motif sequences with physicochemical characteristics similar to SIINFEKL. Such “near neighbor” pentamers were also identified in many other bacteria. This included proteomes of *Listeria monocytogenes*, *Salmonella enterica*, *and Mycobacteria bovis* BCG. *Listeria*, *Salmonella*, and BCG have all been engineered to express OVA similar to the approach shown here with *Brucella* (13, 32, 33). However, no cross-reactivity of native bacterial peptides to OVA specific TCR was reported. This may be because the flanking amino acid context precluded binding and presentation in these bacteria, or that this phenomenon has been observed with other bacteria but not reported (34). The occurrence of near neighbor pentamers in the gastrointestinal microbiome organisms suggests that prior exposure to such peptides is difficult to avoid.

Sequence scans of other *Brucella* species including *B. abortus*, *B. suis*, *B. neotomae*, and *B. ovis* showed these pentamer motif sequences to be conserved. One limitation of this study is that it would not be feasible to delete all genes coding the SIINFEKL near neighbors to definitively confirm the connection between production of SIINFEKL cross reactive peptides and recognition of infected cells by OT-1 T cells. Additionally, the proteins containing the peptides listed in **Table 2** used in these studies may be essential for survival. In searching the literature describing *Brucella* mutants, we could not find engineered mutants of these proteins (35, 36) (37, 38), possibly because they did not survive the mutation process or were severely attenuated.

An intriguing alternative to our explanation could be that *Brucella* infection alters presentation of the host proteome. Indeed, our previous studies have found *Brucella* induces the host Unfolded Protein Response (39, 40) and this ER stress could theoretically alter host or self-antigen presentation. Further, immune activation during *Brucella* infection results in IFN-γ production, which is known to up-regulate MHC I expression and self-antigen presentation. Searching the C57/BL6 mouse proteome revealed no SIINFEKL peptide sequence. However, applying our near neighbor algorithm did reveal 22 instances of ~~~INFEK~ sequence that would potentially be exposed to the T cell. Nevertheless, the predicted binding affinity to H2K^b^ or H2D^b^ was low compared to the *Brucella* peptides used in this study (data not shown). Testing this theory using UPR inducer/inhibitors and a cytokine APC activator did not enhance stimulation of the OVA-specific TCR expressing B3Z T cells. These results suggest the host (mouse) does not present an OVA-like peptide due to ER stress or APC activation by IFN-γ.

Activation of the T cell through the T cell receptor by peptide-MHC I had been thought to be peptide specific; however, our results, along with others (41–45) have shown that T cells can be triggered by peptides with even minimal obvious homology to the primary immunogenic peptide, or amino acid substitutions with similar charge and size. The TCR recognizes an immunogenic complex consisting of peptide bound to MHC I with peptides of 8-11 amino acids in length (46). Of those amino acids, only five are exposed to the TCR (20, 47). Given the genetic combinatorial rearrangement possibilities, an estimated 10^15^ unique TCRs could be generated in the mouse (43). However, studies have shown there are actually <10^8^ distinct TCR clones in the human naïve T cell pool (48), and likely a similar number in mice. Since 20^9^ foreign peptide nonamers can theoretically be generated, it would be mathematically impossible for the T cell pool to recognize all foreign peptides if the TCRs were monospecific (44). Therefore, the TCR must be degenerate and cross-reactive to near-neighbor motifs as demonstrated here.

In fact, a TCR is estimated to react productively with 1 × 10^6^ different MHC-peptides epitopes (49). T cell cross-reactivity has also been documented using the OT-1 transgenic mice model we used in this study (10). Nevertheless, the degeneracy of the TCR correlated with differences in avidity to the peptide-MHC I complexes as others have reported (50). Our studies confirmed that higher concentrations of a degenerate peptide were needed to activate the “SIINFEKL-specific” TCR to the same level activation as SIINFEKL itself. Noteworthy is our observation that antibody to H2K^b^ bound SIINFEKL is apparently not degenerate but SIINFEKL-specific and could only be visualized on *Brucella*-OVA infected cells but not *Brucella* infected cells (Figure 4). Whether this is truly due to antibody specificity or perhaps assay sensitivity would need further investigation.

Our results underscore the complexity and ubiquity of molecular mimicry in T cell recognition. The potential for extensive sharing of nonamers between pathogens, gastrointestinal microbiome and human proteome has been demonstrated, both for MHC I and MHC II (51). This may enable microbes to escape immune surveillance by presenting peptides similar to the host and may also lead to microbial exposure cross-activating autoreactive T cells. Although the association between bacterial infections and autoimmune disorders is still not fully understood (52), recent reports indicate molecular mimicry may be responsible for activation of autoimmune diseases (53–57). Consistent with this, *Brucella* infections have been implicated in several autoimmune diseases (58–61). The genome of *Brucella melitensis* is predicted to encode for 3197 ORFs distributed over two circular chromosomes (62). However, even with this level of complexity, microbe/human commonality is extremely high with 99.7% of human proteins containing bacterial pentapeptides (51, 63). Furthermore, while this study addresses continuous pentamers which are recognized by CD8+ T cells, such peptides are overlaid by the discontinuous pentamers presented by MHC II and recognized by CD4 T cells, with a similar degree of potential cross reactivity (51). Although chicken ovalbumin is not a host protein of mouse or human, we have demonstrated here that there is enough commonality for cross reactivity of several putative *Brucella* peptides with SIINFEKL. It is possible that the cross reactivity of transgenic OT-1 immune cells to *Brucella* could be used in autoimmune studies in mice.

In summary, we have generated a unique tool to dissect CD8+ T cell responses to *Brucella* infection using a widely available TCR transgenic. Further, the OT-1 mice may also be used to probe native *Brucella* infections. Transgenic mice carrying monoclonal T cell receptors are widely used in immunological research. The results presented here raise an important caution for the interpretation of experiments based on reactions to SIINFEKL, or any other small single peptide, unless the presence of the T cell recognition motif or potential cross reactive near neighbors within the host, its microbiome, or an organism under study are addressed. Our results also challenge the assumption that sequence homology will predict molecular mimicry. Thus, using databases comparing sequence of “self” and pathogens will almost certainly underestimate the true contribution of molecular mimicry to pathogen-triggered autoimmunity.

## ACKNOWLEDGMENTS

This work was supported by NIH grants 4R01-AI073558 and F31-AI115931. We thank Dr. J.D. Sauer (UW-Madison) for his gift of B3Z cells and pPL2erm-ActA100-B8R-OVA and Dr. M. Suresh (UW-Madison) for his gift of DC2.4 cells. We thank Dr. M. Imboden (ioGenetics) for insight and discussion during the project. The authors have no conflict of interest to declare. EJH and RDB are equity holders in ioGenetics LLC.

